# Cell Lineage-Guided Microanalytical Mass Spectrometry Reveals Increased Energy Metabolism and Reactive Oxygen Species in the Vertebrate Organizer

**DOI:** 10.1101/2023.07.07.548174

**Authors:** Aparna B. Baxi, Jie Li, Vi M. Quach, Peter Nemes

## Abstract

Molecular understanding of the vertebrate Organizer, a tissue center critical for inductive signaling during gastrulation, has so far been limited to transcripts and some proteins due to limitations in detection and sensitivity. The Spemann-Mangold Organizer (SMO) in the South African Clawed Frog (*X. laevis*), a popular model of development, has long been discovered to induce the patterning of the central nervous system. Molecular screens on the tissue have identified several genes, such as goosecoid, chordin, and noggin, with independent ability to establish a body axis. A comprehensive study of proteins and metabolites produced in the SMO and their functional roles has been lacking. Here, we pioneer a deep discovery proteomic and targeted metabolomic screen of the SMO in comparison to the rest of the embryo using liquid chromatography high-resolution mass spectrometry (HRMS). Quantification of ∼4,600 proteins and a panel of metabolites documented differential expression for ∼450 proteins and multiple intermediates of energy metabolism in the SMO. Upregulation of oxidative phosphorylation (OXPHOS) and redox regulatory proteins gave rise to elevated oxidative stress and an accumulation of reactive oxygen species in the Organizer. Imaging experiments corroborated these findings, discovering enrichment of hydrogen peroxide in the SMO tissue. Chemical perturbation of the redox gradient affected mesoderm involution during early tissue movements of gastrulation. HRMS expands the bioanalytical toolbox of cell and developmental biology, providing previously unavailable information on molecular classes to challenge and refine our classical understanding of the Organizer and its function during early patterning of the embryo.

## INTRODUCTION

During early vertebrate development, the processes of germ layer formation, body patterning, and cell differentiation rely on morphogen gradients that are generated from signaling centers within the embryo. One such tissue is the vertebrate Organizer, referred to as the Node in human, chick, and mouse, the Shield in zebrafish, and the Spemann-Mangold Organizer (SMO) in amphibians (1, 2). The Organizer transiently forms on the dorsal side of the embryo and orchestrates dorsal-ventral patterning of the mesoderm, induction of the neural ectoderm, and morphogenetic movements during gastrulation. Classical embryological experiments in many vertebrate species have demonstrated the inductive prowess of the Organizer, wherein transplantation of this tissue to the ventral side of an embryo resulted in the formation of a secondary body axis containing properly patterned mesoderm and induced neural ectoderm (3-6). This surprising finding has kindled tremendous interest in uncovering the molecular composition of the Organizer, which so far has been studied primarily through expression cloning and transcriptomic screens (7-11). However, our understanding about the proteome and metabolome, the comprehensive suite of proteins and metabolites that carry out molecular functions, including signaling, is still lacking in the Organizer for any animal.

The South African clawed frog (*X. laevis*) provides a tractable and scalable model to decipher the SMO. The SMO secretes several diffusible molecules (morphogens) that signal surrounding cells to begin the formation of specialized tissue (2, 12). Numerous studies based on expression cloning have discovered molecules secreted by the SMO using *X. laevis* (6, 7, 13-15), where embryonic development is fast and external to the mother, yielding ready access to microsurgery at the gastrula stage. Expression cloning and cDNA expression libraries constructed from the SMO have identified morphogens, such as secreted anti-BMP factors (e.g., Chordin, Noggin, Xnr3, Follistatin) and secreted anti-Wnt factors (e.g., Dkk, Frzb-1, Cerebreus), to be critical for patterning the mesoderm and inducing the neural ectoderm (6, 13, 16-19). However, these technologies have been limited to targeting one or a few gene candidates at a time. Recent transcriptomic screens have uncovered additional morphogens, revealing Pkdcc1 and Bighead, which support head formation by antagonizing Wnt signaling (10, 20). Therefore, investigation of gene products in the *X. laevis* SMO is vital for expanding our basic understanding of the vertebrate Organizer.

Our knowledge of proteins and metabolites produced downstream of transcription in the SMO remains elusive, primarily due to technological limitations in detection, quantification, and sensitivity. Previous studies found that transcript abundances do not necessarily predict the abundance of proteins, especially in dynamic systems such as the developing *Xenopus* embryo (21, 22). Recent advances in high-resolution mass spectrometry (HRMS) by us and others enabled the identification of thousands of different proteins in *X. laevis* embryonic tissues (23-25) and hundreds of metabolites (26, 27) in a discovery setting, without necessitating functioning probes such as antibodies for detection. Technical advances in microanalytical sampling, processing, separation, and ionization have recently primed HRMS to the point of being able to uncover spatial and temporal proteomic (28-31) and metabolomic (27, 32) differences between identified embryonic cells (blastomeres) within 8-to 128-cell *Xenopus* embryos, leading to the discovery of metabolites capable of altering early cell-fate decisions (27, 33). Therefore, advanced HRMS and *Xenopus* present a unique integration and potential to make leaps forward in our knowledge of proteins and metabolites and their function during vertebrate development.

Here, we sought to perform a comprehensive molecular screen to identify proteome and metabolome signals in the SMO. We utilized embryological microinjection to label the Organizer for microsurgical isolation. Using liquid chromatography HRMS, we quantified ∼4,600 proteins, including ∼450 proteins with differential regulation between the SMO and the rest of the embryo (RE). Detection of germ layer specific transcription factors such as Brachyury (Tbxt) and Sox3 indicated sufficient sensitivity to appreciate molecular mechanisms in the SMO at the level of proteins. Our results revealed an enrichment of oxidative phosphorylation and energy metabolism in the SMO. Targeted HRMS assays on the metabolite intermediates produced downstream, complemented by classical fluorescence-based metabolite assays when available, revealed local oxidative stress and enrichment of reactive oxygen species (ROS) in the tissue. Through chemical perturbation, we tested the implications of this redox gradient on regulating cellular involution at the blastopore lip during gastrulation. These previously unknown molecules highlight the importance of revisiting key developmental events through a multi-omics lens to gain a holistic view of the bimolecular processes orchestrating embryonic development.

## RESULTS

### Lineage-Guided Dissection of the Organizer

Our experimental approach (**Fig. 1**) began with isolation of the SMO tissue and the remainder of the embryo (RE). Inspired by the classical expression cloning approaches (10, 13, 34), we identified the precursor cells in the 32-cell stage embryo that give rise to stereotypical tissue lineages (35, 36) with expression of known SMO-specific genes, such as *goosecoid* and *chordin* (37). **Figure 1** presents two-color, dual fluorescent labeling of the dorsal-animal cells (D112 in red) and the dorsal-vegetal cells (D212 in green) and their respective descendent clones in the gastrula (stage 10). We considered injection of the left and right D112 and 212 cells (4 cells per embryo) to sufficiently label the SMO in this study (**Fig. 1A**). The fluorescently labeled tissue was dissected using a pair of sharpened forceps (**Fig. 1B**) and the remainder embryo (RE) was also collected to serve as a reference. The isolated tissues were processed for discovery proteomic and targeted metabolomic profiling (**Fig. 1C**). The proteome digests were barcoded with isobaric mass tags (126 to 131 tandem mass tags) to permit multiplexing relative quantification via nanoLC quadrupole–ion trap–orbitrap tribrid HRMS executing simultaneous precursor selection tandem MS (MS^2^) for identification and quantification at the 3^rd^ order (MS^3^) (**SI *Methods***). A panel of selected metabolites (below) were absolutely quantified using external concentration calibration via LC quadrupole–time of flight MS^2^ (***Methods***).

**Figure 1.**
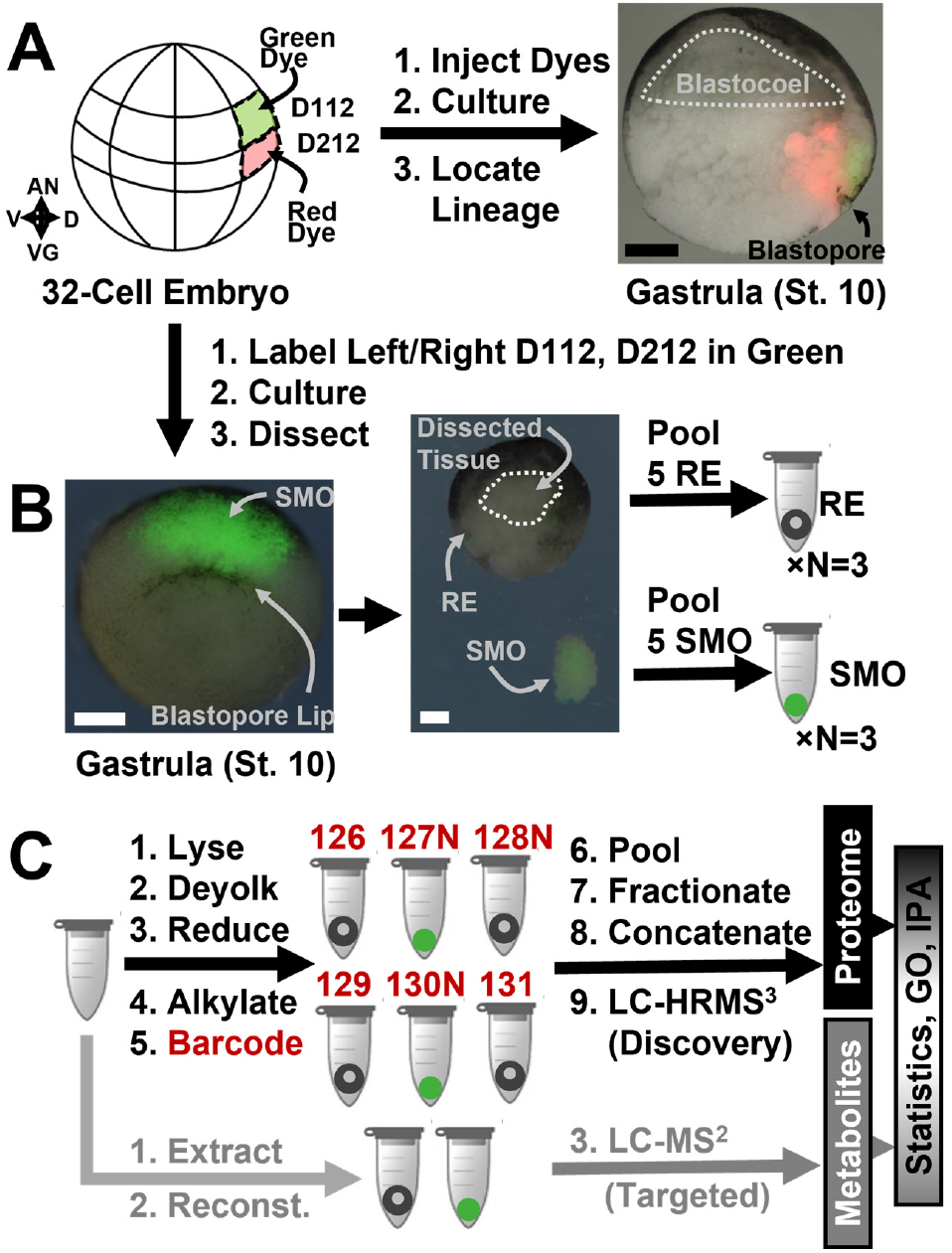
Workflow for lineage tracing and dual proteomic-metabolomic screening of the Spemann-Mangold Organizer (SMO) in *X. laevis* and high-resolution mass spectrometry (HRMS). **(A)** The dorsal midline equatorial cells D112 and D212 of the 32-cell embryo (**left panel**) were lineage traced to encompass the majority of SMO tissue, shown in the hemisected gastrula at stage 10 (**right panel**). **(B)** The lineage-traced tissues were dissected to pool five SMO tissues and the remainder of the embryo (RE) as one biological replicate. **(C)** Five different pools of tissues were processed for tandem/multistage HRMS quantification of the proteome via isobaric barcoding bottom-up proteomics (MS^3^) and metabolite intermediates via targeted metabolomics (MS^2^). The resulting proteomics-metabolomics dataset was statistically analyzed and interpreted using gene ontology and molecular pathway analysis. Scale bars, 250 μm.

### Proteomic Profiling of the Organizer

We identified ∼4,600 proteins in the tissues through bottom-up proteomics using HRMS. The technical aspects of the analysis are detailed in the electronic **Supplementary Information** (**SI**) document. As shown in **Figure 2A**, about ∼450 proteins were significantly differently enriched between the SMO and RE (*p* <0.05, Student’s t-test, **SI Table 1A**). Approximately 160 proteins were more abundant in the SMO, whereas ∼300 proteins had higher levels in the RE. We inquired about the canonical roles of these proteins using ingenuity pathway analysis (IPA). **Figure 2B** represents the 15 pathways with the highest and lowest z-score in the SMO, a metric of significance for differential regulation. The upregulated pathways included proteins involved in cellular energy metabolism such as oxidative phosphorylation (OxPhos), the TCA cycle, and fatty acid oxidation. The data also indicated an enrichment of processes pertaining to translation initiation, regulation, and inhibition of mRNA degradation (eukaryotic translation initiation factors, such as Eif, Eif1g, and Eif1ha). Several signaling pathways were downregulated controlling cell adhesion, migration, and motility (e.g., ephrin, integrin-linked kinase or ILK, mTOR, and paxillin) and proliferation (chemokine cell receptor CXCR4 and interleukin-8, IL-8). These results suggest that cells in the SMO elevate protein synthesis and metabolic energy production but reduce motility and division immediately before gastrulation (stage 10).

**Figure 2.**
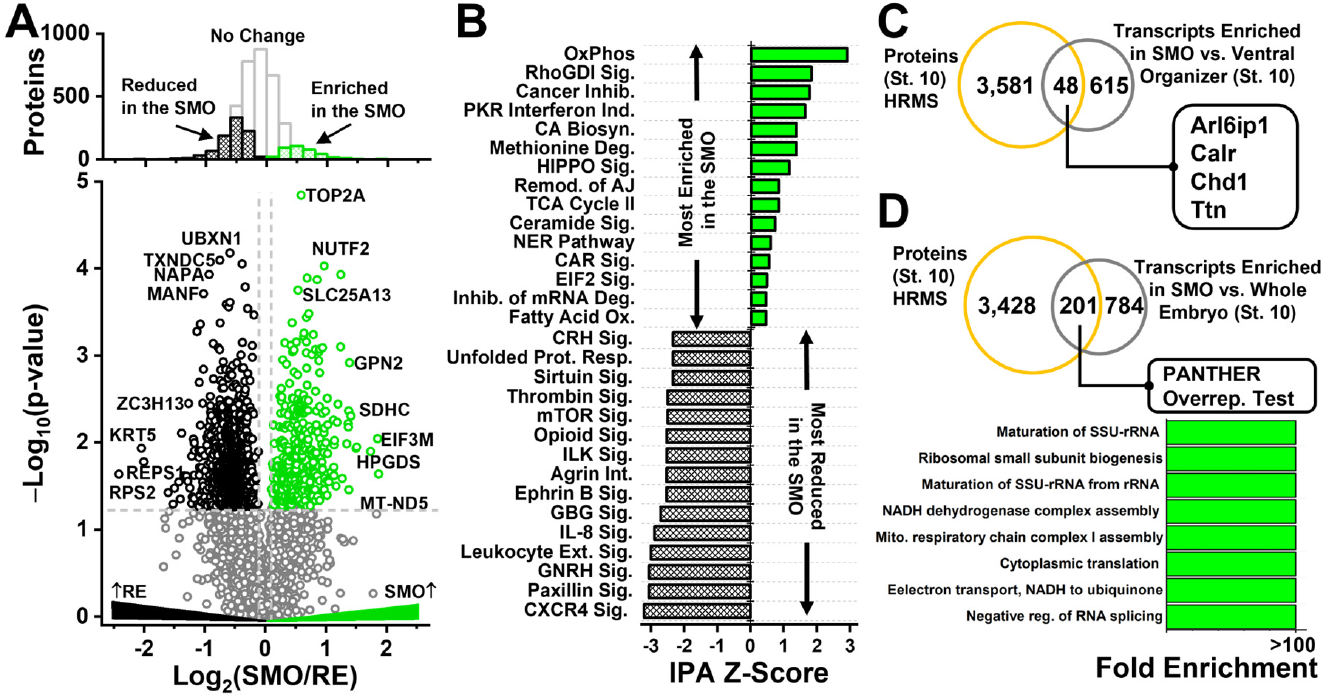
Proteomic profiling of the Spemann-Mangold Organizer (SMO) vs. the remainder embryo (RE). (**A**) Of 4,600 proteins identified, ∼450 showed differential enrichment in either the SMO (green) or the RE (black). (**B)** Functional annotation of the 15 canonical pathways with most significant z-scores via Ingenuity Pathway Analysis. Protein upregulation in the SMO from HRMS on the SMO in comparison to **(C)** elevated gene expression vs. the dorsal organizer from Ref. (10). and **(D)** single-cell RNA-sequencing from Ref. (38). (Inset) Analysis of biological pathway overrepresentation for gene products enriched both based on HRMS and single-cell RNA-seq using PantherDB with Bonferroni correction for multiple testing.

These proteomics results complemented those available from transcriptomics screens, also at the gastrula. Transcriptome profiling previously revealed ∼650 transcripts enriched in the SMO in comparison to the ventral organizer (10). From this dataset, 48 genes intersected with our HRMS-based deep proteome analysis; the proteins translated from 4 of these genes we quantified also with enrichment in the SMO (**Fig. 2C**). Single-cell sequencing (Seq) independently found ∼1,000 genes to be differentially transcribed (38) in the Organizer. Compared to this dataset, 201 genes overlapped with our proteome data, only 39 of which yielded elevated protein levels in the SMO based on our HRMS quantification.

Overrepresentation analysis in PANTHER (**Fig. 2D**) concluded upregulation of various mitochondrial functions related to energy production (e.g., electron transport, energy production and utilization, respiration, and protein translation).

### Canonical Components of Energy Production

The HRMS data offered a vantage point to appreciate several pathways of energy metabolism, multiple at once. **Figure 3** profiles the abundance of known cytoplasmic proteome components of glycolysis and the pentose phosphate pathway (PPP) as well as OxPhos and the Krebs (TCA) cycle. Quantification of proteins in these specific pathways are shown in **Figure S1**. While most components of glycolysis–PPP were comparable in abundance across the tissues, downregulation of pyruvate kinase M (Pkm) and one isoform of glyceraldehyde-3-phosphate dehydrogenase (Gapdh) created bottlenecks in the pathway. Pyruvate dehydrogenase (e.g., pyruvate dehydrogenase E1α, or Pdha), which links glycolysis to the TCA cycle, was also similar between the SMO and RE. In stark contrast, key protein components of nearly all complexes of mitochondrial electron transport chain and the TCA cycle were significantly upregulated in the SMO. Therefore, we proposed that SMO was a site of active energy production through mitochondrial function (OxPhos), rather than the alternative cytoplasmic sources.

**Figure 3.**
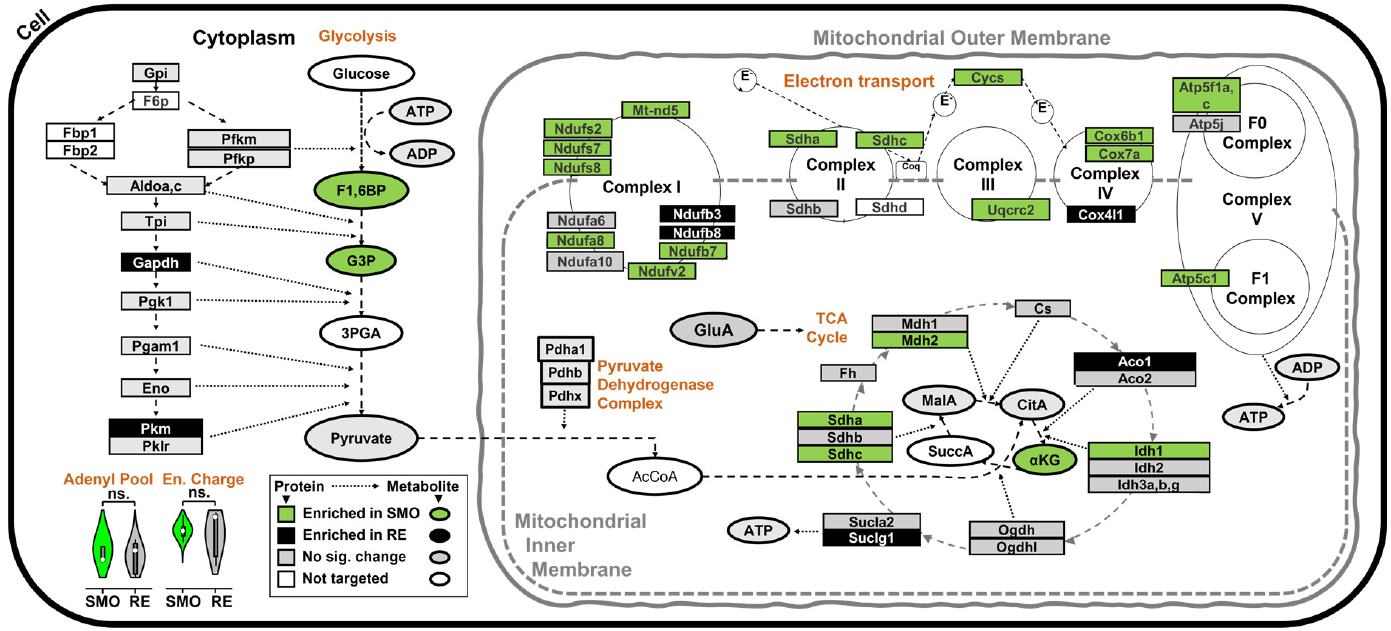
Proteome and targeted metabolite profiling of energy production between the SMO and RE using HRMS. Protein downregulation in glycolysis and upregulation in oxidative phosphorylation and TCA intermediates established enrichments for several metabolite intermediates in the SMO in comparison to the RE tissue. Key to metabolite labels: CitA, citric acid; F1,6BP, fructose-1,6-biphospate; G3P, glyceraldehyde-3-phospate; MalA, malic acid; α-KG, α-ketoglutaric acid.

This hypothesis was testable through the lens of HRMS into the metabolome. We developed targeted LC-MS assays for a panel of metabolites to quantify their tissue-endogenous concentrations. The technical details of the analysis are provided in the **SI** document. Specifically, external linear calibration (***Methods*, Fig. S2**) aided us in quantifying fucose-1,6-biphosphate (F1,6BP), glyceraldehyde-3-phosphate (G3P), and pyruvate from glycolysis– PPP as well as α-ketoglutaric acid (α-KG), citric acid (CitA), glutamic acid (GluA), and malic acid (MalA) from the TCA cycle. The overall state of tissue energy was assessed through its metabolic currencies adenosine triphosphate (ATP), adenosine diphosphate (ADP), and adenosine monophosphate (AMP). Enrichment of F1,6BP and G3P in the SMO and comparable pyruvate production (data in **Fig. S3A**) supported the likely presence of a glycolytic enzymatic bottleneck under control by downregulation of the Gapdh protein in the SMO.

The analysis also explored mitochondrial energy production. As illustrated in **Figure 3**, multiple components of Complexes I, II, III, IV, and V of the electron transport chain were enriched in the SMO. These data would suggest enhanced ATP production, but our adenylate assays on ATP, ADP, and AMP confirmed similar activities between the SMO and RE. The adenylate energy pool, which we quantified using the popular metrics of adenylate energy charge and [ATP]/[ADP] ratios (*Methods*) of ∼0.97 and ∼35, respectively (**Fig. S3B, *Methods***). This ATP/ADP concentration ratio is considerably high, even among subcellular organelles, indicating oxidative, rather than glycolytic energy source for the SMO.

For the TCA cycle, where multiple proteins were also upregulated, all targeted products were comparable between the tissues (data in **Fig. S3C**), except for α-ketoglutarate (α-KG) that was produced in ∼3-times higher concentration in the SMO compared to the RE. Production of this compound would align with recent studies linking α-KG to moonlighting roles in signaling, regulation of gene expression via DNA demethylation, and redox homeostasis in addition to its more prominent function in generation of glutamate and glutamine for protein synthesis (39-41). Our assays revealed comparable glutamic acid levels (**Fig. S3C**). While rapid energy production aligns with active protein translation and metabolism, signaling is a well-established function of the SMO.

### Elevated Reactive Oxygen Species in the SMO

An upregulation of fatty acid oxidation (increased Eci1 expression) and OxPhos in the SMO, as found from our proteomic analysis (**Fig. 2B**), suggested also elevated production of OxPhos-generated reactive oxygen species in the SMO. To assess the spatial localization of ROS in the embryo, we incubated live, stage-9 embryos (before gastrulation begins) in a probe that fluoresces upon reaction with cellular H_2_O_2_ (**Fig. 4, SI *Methods***), followed by culturing to stage 10 when our SMO studies were performed. Live fluorescence imaging of embryos revealed local enrichment of H_2_O_2_ in the dorsal equatorial/vegetal region of the embryo. Specifically, the fluorescence from the H_2_O_2_-sensitive probe was maximal above the blastopore lip the region (**Fig. 4A**), with its intensity tapering down toward the dorsal equatorial region (**Fig. 4B**). Fluorescence from the probe was minimal in other regions of the embryo, such as the animal pole (**Fig. 4C)**. In a systematic approach, we quantified regional fluorescence intensities in the SMO and the vegetal-side region non-SMO regions defined in **Fig. 4C**) among N = 8 embryos (**SI *Methods***). Statistics on the data indicated significant H_2_O_2_ enrichment in the SMO region.

**Figure 4.**
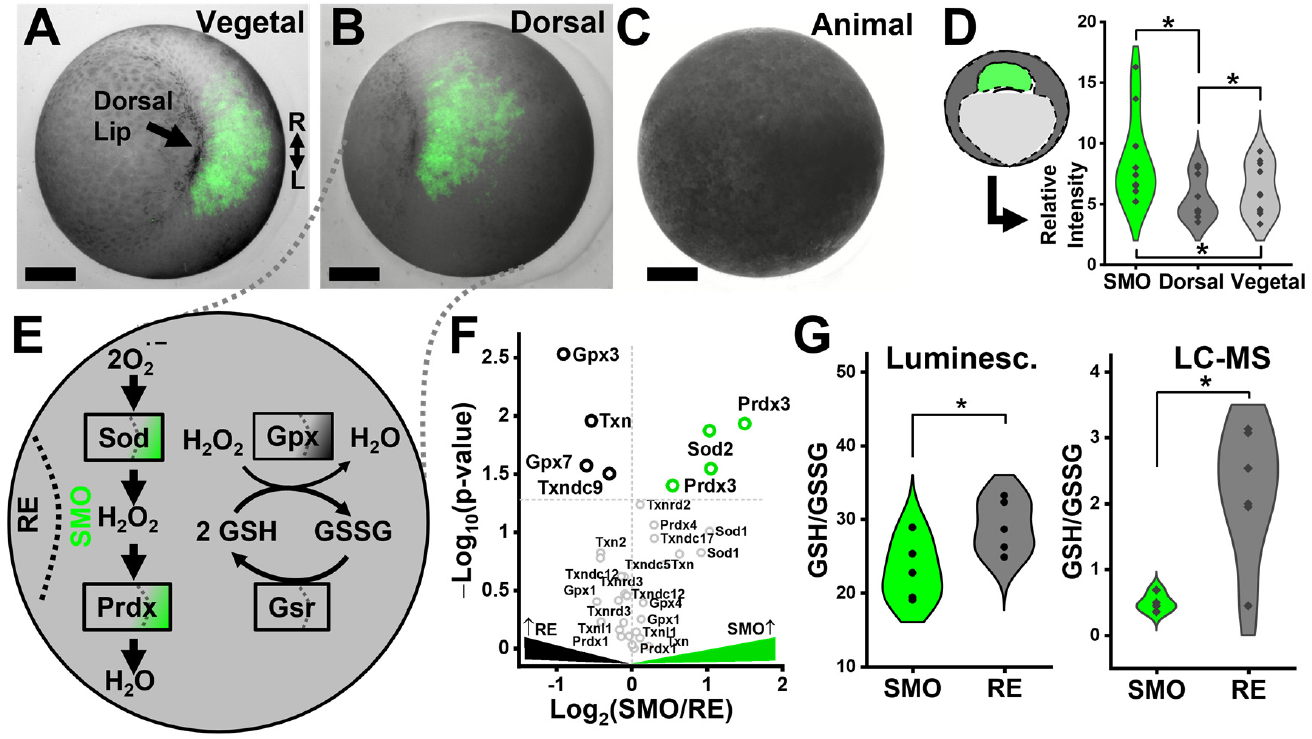
Our biochemical model of elevated oxidative stress in the SMO region. H_2_O_2_-sensitive fluorescent imaging of the **(A)** vegetal, **(B)** dorsal, and **(C)** animal poles of stage-10 *X. laevis* embryos. **(D)** Background-normalized fluorescence intensity for BioTracker green H_2_O_2_ live cell dye measured in the SMO, dorsal, and vegetal regions of the embryo. Key: \**p* < 0.05 (paired Wilcoxon signed ranks test). **(E)** Our working molecular model proposing active generation of H_2_O_2_ based on enrichment of the redox proteins Sod, Prdx, and Gpx and downregulation of Gpx, a protein converting glutathione (GSH) to oxidized glutathione (GSSG), the buffers of the redox system. **(F)** HRMS testing of the hypothesis through quantification of the redox system proteome. Key: \**p* < 0.05 (Student’s t-test). **(G)** Experimental profiling of GSH/GSSG using a luminescence-based assay **(left panel)** and targeted LC-MS metabolomics **(right panel)** validated elevated redox stress. Key: \**p* < 0.05 (Student’s t-test). Scale bars, 250 μm.

Our developing molecular model was honing on oxidative signaling and stress (**Fig. 4E**). Based on the measured upregulation in redox processes (**Fig. 2**) and the many mitochondrial complexes in the electron transport chain (**Fig. 3**), we posited a potential presence for reactive oxygen species. **Figure 4F** quantifies underlying key proteins, such as superoxide dismutases (Sod), peroxiredoxins (Prdx), thioredoxins (Trx), and glutathione peroxidases (Gpx). The Sods carry out dismutation of superoxide radicals generated as a by-product of oxygen metabolism into molecular oxygen and H_2_O_2_ (**Fig. 4E**). Cytosolic Sod1 was comparable between the tissues, and Sod2 was substantially upregulated in the SMO (**Fig. 4F**), which would lead to elevated H_2_O_2_ production. While isoforms of Prdx3 were elevated in the SMO, H_2_O_2_ reduction is catalyzed by Prdx and facilitated via GSH→GSSG oxidation under catalysis by Gpx (**Fig. 4E**). Isoforms of Gpx were not significantly different between the SMO and the RE, with an exception to GPX3 and 7, which were enriched in the RE (**Fig. 4F**). Therefore, this model predicted the observed H_2_O_2_ pool in the SMO.

Our model also predicted potential differences in redox state and a resulting oxidative stress (**Fig. 4E**). To put cellular oxidative stress on trial, we characterized the glutathione (GSH)–oxidized glutathione (GSSG) pool in the tissues. A luminescence assay found reduced GSH/GSSH ratios in the SMO (**Fig. 4G, *Methods***). To enhance the molecular specificity of detection and sensitivity of quantification, we also developed an LC-MS assays for quantifying GSH/GSSG (***Methods***). This targeted metabolomic MS method confirmed decreased GSH/GSSG in the SMO compared to RE (**Fig. 4G**). Therefore, the SMO is a site of elevated ROS (H_2_O_2_) production and oxidative state in the gastrula.

### Depletion of ROS Causes Defects in Gastrulation Movement

Increased ROS production in the SMO may fulfill a function, we put forward. As illustrated in the experimental design in **Figure 5A**, we pursued chemically depleting the ROS in the SMO to monitor gastrulation movements between stages 10 and 11. In N = 10 embryos at stage 10, the SMO was injected with the antioxidant agent N-acetyl cysteine (NAC), which is known to buffer ROS by upregulating glutathione production. In parallel, sibling embryos were injected with 20% Steinberg’s solution (control), the embryo culture media. A reduction in fluorescence from the H_2_O_2_-reactive dye confirmed decreased ROS production following NAC injection in the SMO in N = 10 embryos.

**Figure 5.**
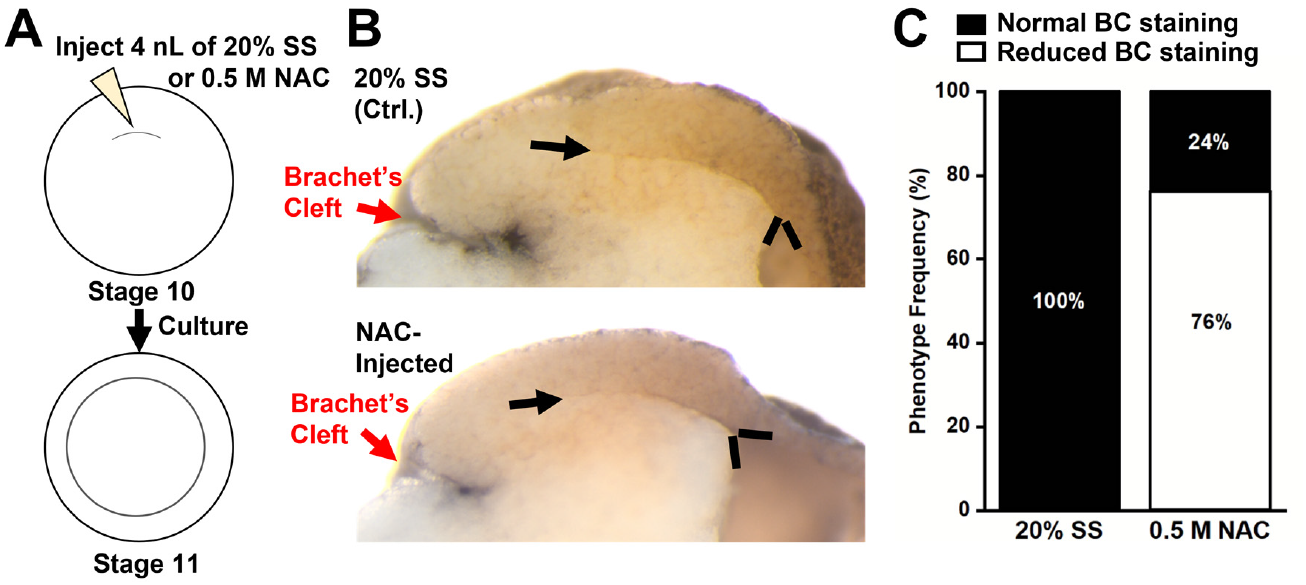
Functional assessment of ROS enrichment in the SMO on mesoderm involution during gastrulation. **(A)** The SMO region of stage-10 embryos was injected with the antioxidant N-acetyl cystine (NAC) or control 20% Steinberg’s solution (SS) and cultured to stage 11. **(B)** Fibronectin immunostaining at stage 11 revealed deeper involution of Brachet’s cleft (red arrow). Black arrows show the involuted mesoderm. **(C)** Validation of the phenotypes based on fibronectin staining in the cleft of NAC-injected embryos compared to the control siblings.

Reduced ROS levels affected tissue involution during gastrulation. We performed immunostaining for fibronectin (**Fig. 5B**) for the 10 embryos in each group, as mesoderm involution requires fibronectin (FN) deposition along Brachet’s cleft (42, 43). The analysis revealed NAC injection to delay involution of the mesoderm by mid-gastrulation (stage 11), with ∼76% of the experimental embryos showing pronouncedly decreased fibronectin staining along the cleft vs. the control siblings (**Fig. 5C**). The cleft was open rather than tightly opposed to the involuted mesoderm. Gene expression analysis via *in situ* hybridization was positive for the neural markers *foxd4* and *sox2* and mesodermal marker *chordin* in NAC-injected embryos (data not shown), indicating that the embryo eventually compensated for the degree of ROS depletion that was experienced with during gastrulation in this study.

## DISCUSSION

This study strategically integrated the frog embryo as a vertebrate developmental model and HRMS as a quantitative multi-’omics analyzer to expand our historically gene- and transcript-centric understanding of the Organizer. Using HRMS, we drafted the SMO proteome through simultaneous detection of ∼4,600 proteins. Apprximately 450 of these proteins were differentially a‘bundant between the SMO and RE. The data paint a molecular model of upregulated mitochondrial energy production. Reduction in enzyme levels forms a bottleneck to accumulating sugar intermediates via glycolysis–PPP, but not pyruvate as feed to the energy demand. Likewise, comparable adenosine ATP, ADP, and AMP pools keep regular energy reservoirs. An upregulated OxPhos gives rise to active mitochondrial electron transport, producing ROS, including the surprising accumulation of H_2_O_2_ and α-KG through an upregulated TCA cycle. Elevated GSH/GSSG buffer suggested elevated oxidative stress, raising a potential noncanonical role as morphogens embryonic patterning.

These findings add to a body of growing evidence on the metabolic patterning of the embryo. Recent work has begun to highlight the significance of ROS-dependent oxidative “eustress” in the context of stem cell differentiation and neurulation (44, 45). These experiments demonstrated a regulatory role of ROS in driving thiol-oxidation based modulation of signaling proteins to regulate development (46), including regulation of Wnt/β-catenin in a duration- and context-dependent manner. Treatment with a low dose of H_2_O_2_ stabilized β-catenin leading to a concomitant increase in the expression of endogenous Wnt target genes On the other hand, exposure to exogenous H_2_O_2_ has also been shown to negatively modulate the Wnt/β-catenin pathway by decreasing the amount of β-catenin that is localized in the nucleus (47). The SMO transcriptome is shown to be induced by early β-catenin signaling (48) and knocking down β-catenin inhibits the formation of a dorsal axis.

Therefore, our HRMS studies suggest a noncanonical potential for ROS-based regulation to nuclear localization of β-catenin in the SMO. Indeed, depletion of ROS with the use of antioxidant (NAC) detectably disrupted the involution of the mesoderm during gastrulation while maintaining tissue specification. The precise mechanisms though which ROS play a role in regulating functions of the SMO remains to be the focus of follow-up work. Additionally, the discovery-proteomics and metabolite-targeted HRMS assays yielded previously unavailable information on the molecular component of the embryo, which can be used to test existing and generating hypotheses regarding the function of vertebrate Organizer.

## METHODS

Comprehensive descriptions of technologies and methods are provided in the **SI** document. All protocols ensuring the humane treatment of animals were approved by the Institutional Animal Care and Use Committee (approval no. R-FEB-21-07) of the University of Maryland, College Park.

## AUTHOR CONTRIBUTIONS

P.N. and A.B.B. designed the study; A.B.B. and V.M.Q. dissected the Spemann-Mangold organizer. A.B.B. processed and measured the proteins.

A.B.B. performed the optical imaging. J.L. developed and performed metabolomics assays.

A.B.B, J.L., and P.N. analyzed the data. A.B.B. and P.N. interpreted the results and prepared the publication. All co-authors have commented on the report.

## ACKNOWLEDGEMENT

This work was supported by the National Institutes of Health (award no. 1R35GM124755 to P.N.). We thank Dr. Sally Moody (George Washington University, Washington, DC) for providing reagents and instrumentation for *in situ hybridization* and antibody staining. We thank Leena Pade (University of Maryland, College Park) for assistance during preparation of the manuscript.

## Data Repository

The MS proteomics data were deposited to the ProteomeXchange Consortium via the PRIDE partner repository with the DOI dataset identifier PXD043583. For MS metabolomics data, the data for this study will be accessible at the NIH Common Fund’s NMDR website (supported by NIH award no. U01-DK097430), the Metabolomics Workbench, https://www.metabolomicsworkbench.org.

