## Supplementary material for "Cell Lineage-Guided Microanalytical Mass Spectrometry Reveals Increased Energy Metabolism and Reactive Oxygen Species in the Vertebrate Organizer": SI Document

### MATERIALS AND METHODS

**Materials and Reagents.** Unless described otherwise, liquid chromatography (LC) mass spectrometry (MS)-grade solvents, reagent-grade chemicals, MS-grade trypsin, and Alexa Fluor 488 (10,000 g/mol MW, anionic fixable, green fluorescent dextran) were purchased from Fisher Scientific (Pittsburgh, PA). Cysteine, trizma (Tris) hydrochloride, trizma base, combretastatin A-4, cytochalasin D, and nonidet P-40 substitute (NP-40) were from Sigma-Aldrich (St. Louis, MO).

**Solutions.** For culturing embryos, 100% and 50% (w/v) Steinberg's solutions (SS) were prepared as described elsewhere (1). Tissues were fixed in 4% paraformaldehyde in phosphate buffered saline following standard protocols (2). To deplete tissues of yolk, lysis was performed in 250 mM sucrose, 1% NP-40, 5 mM EDTA, 20 mM Tris-HCl, 10 µM combretastatin 4A, and 10 µM cytochalasin D. The *metabolite extraction solvent* was prepared to contain 40% (v/v) acetonitrile (ACN) and 40% (v/v) methanol (MeOH) in water, cooled to 4 °C. The *protein extraction/digestion buffer* was prepared to contain 50 mM ammonium bicarbonate (AmBic) in water.

**Animal Care and Embryology.** Adult male and female *Xenopus laevis* were purchased from Nasco (Fort Atkinson, WI). The frogs were maintained in a breeding colony and handled following protocols approved by the Institutional Animal Care and Use Committee (approval no. R-FEB-21-07) of the University of Maryland, College Park. Standard protocols were followed to obtain embryos using *in vitro* fertilization (1). Embryos were dejellied in 2% (w/v) cysteine solution (pH 8) and cultured in 100% SS. Embryonic development was staged following the guidelines established by Nieuwkoop and Faber (NF) (3). To ensure accurate cell fate mapping, this study only used 32-cell embryos (NF stage 6) that developed from 2-cell embryos (NF stage 2) with stereotypical pigmentation and cleavage across the dorsal-ventral axis (4). Cell clones of the SMO were traced by microinjecting of their precursor D112 and D212 cells (**Fig. 1**) with 1 nL of 0.5% Alexa Fluor 488 (10,000 g/mol MW) using a micromanipulator-operated micropipette connected to a microinjector (PLI 100A, Warner Instruments, Hamden, CT). The labeled embryos were raised in 50% SS to early gastrula (NF stage 10). Live development was monitored with a stereomicroscope (SMZ18, Nikon Instruments Inc., Melville, NY).

**Microsurgical Tissue Dissections.** The SMO was lineage-traced under epifluorescence and isolated in a 2% (w/v) agarose-coated Petri dish containing 50% SS. The dissected SMO tissues and the remainder embryo (RE) were collected in LoBind Eppendorf tubes separately, and processed using our established protocols for proteomic (5-7) or metabolomic (7-9) analysis. <u>For proteomics</u>, a total of 5 dissections for each biological replicate (BR) sample were collected and frozen on dry ice. The tissues were stored at −80 °C until further processing. In total, 3 independent BRs were collected. <u>For metabolomics</u>, complementary detection technologies were utilized. For Glo assays (Promega, WI), 3 sets of dissected SMO and RE tissues were pooled as 1 BR and stored at −80 °C until analysis. For MS-based metabolomics of the TCA cycle and glycolysis, 10 dissections of SMO and RE were pooled as 1 BR. For targeted quantification of reduced and oxidized glutathione, 20 tissues of SMO and RE were dissected and pooled as one BR. The dissected tissues were stored in LC-MS-grade methanol at −80 °C until sample processing.

**Sample Preparation for Quantitative Proteomics.** The dissected tissues were lysed in 200 µL of yolk depletion buffer for the SMO and 800 µL for the RE, to account for proportionally larger amounts of proteins available in the latter. The resulting samples were incubated at 4°C for 10 min and centrifuged at 2,500 × *g* at 4 °C for 4 min to pellet yolk platelets. The supernatant containing the deyolked lysate was separated from the pellet by gentle pipetting. The RE lysate was halved before further processing. The proteome was denatured by addition of SDS to 1% (v/v) final concentration, reduced using 5 µL (SMO samples) and 10 µL (RE samples) of 0.5 M dithiothreitol with incubation at 60 °C for 30 min. The proteins were alkylated using 15 µL of 0.5 M iodoacetamide in the dark at room temperature for 20 min, before quenching the reaction with 5 µL of dithiothreitol. The processed proteome was purified by precipitation in cold acetone (overnight, –20 °C). The resulting protein pellet was suspended in 50 mM AmBic and digested with 1 µg trypsin (~1:50 protein: protease ratio) overnight at 37 °C. Proteins from the RE were processed similarly using double amounts of reagents to account for larger tissue amounts. The total amount of resulting peptides was estimated based on absorbance at 205 nm in 50 mM triethylammonium bicarbonate (Synergy HTX, BioTek, Winooski, VT).

To enable multiplexed protein quantification, equal amounts of total peptides from each of N = 3 biological replicates were tagged with tandem mass tags (TMT10plex using 126, 127N, 128N, 129N, 130N, and 131 channels, lot QK223163, Thermo Fisher Scientific, Waltham, MA). A total of ~100 µg of TMT-labeled peptide mixture was fractionated under high pH in spin columns following vendor recommendations (Pierce High pH Reversed-Phase Peptide Fractionation Kit, Thermo). The peptides were eluted in 0.1% (v/v) trimethylamine in water containing ACN at 10%, 12.5%, 15%, 17.5%, 20%, 22.5%, 30%, and 50% (v/v). Each resulting fraction was dried in a vacuum concentrator (Labconco, Kansas City, MO) and stored at –80 °C. The peptide samples were thawed and dissolved in 5% ACN containing 0.1% (v/v) formic acid (FA) for analysis by nano-flow LC (nanoLC) MS.

**Sample Preparation for Quantitative Metabolomics.** The tissues frozen in methanol were defrosted and vacuum-dried (Labconco) at 4 °C. The metabolome was extracted in a 500 µL mixture of 40% (v/v) ACN and 40% (v/v) methanol in water. Samples were vortexed for 30 s at room temperature, sonicated for 10 min in an ice-cold bath, followed by a 10 min incubation in the ice-cold bath. This process was repeated twice. After metabolite extraction, the samples were incubated in a –20 °C freezer for 1 h, followed by vortexing for 30 s and centrifugation for 10 min at 4 °C and 13,000 × *g* to pellet insoluble cell components. The supernatant was transferred to a new 1.5 mL Eppendorf vial, dried at 20 °C, and reconstituted in 45 μL of LC-MS grade water. For GSH/GSSG analysis, the supernatant was reconstituted in 45 μL of 50% ACN. The supernatant was centrifuged at 13,000 × *g* for 10 min at 4 °C, then stored at –80 °C until LC-MS measurement.

**Nano-LC MS Proteomics**. A total of 1 µg of peptide mixture was separated on a C18 column (75 µm inner diameter, 2 µm particle size with 100 Å pores, 50 cm length, Acclaim PepMap 100, Thermo). Separation was performed at 300 nL/min in a nano-LC system (Dionex Ultimate 3000 RSLCnano, Thermo) providing a 150-min gradient between Buffer A (100% water, 0.1% FA) and Buffer B (100% ACN, 0.1% FA). The sample was loaded in 95% Buffer A containing 5% Buffer B and eluted via a multistep gradient that ramped Buffer B as follows: held at 5% for 5 min, then ramped to 10% over 15 min, to 26.5% over 150 min, to 80% over 152 min, held at 80% for 3 min. To recover and condition the analytical column for the next injection, the mobile phase was then ramped to 5% over 5 min, then the solvent composition was held for 15 min.

Peptide ions were detected in an orbitrap-quadrupole-ion trap tribrid high resolution mass spectrometer (Orbitrap Fusion Lumos, Thermo). Peptides were ionized in a nanoelectrospray ionization (ESI) source consisting of pulled fused silica capillary (10 µm tip, New Objective, Woburn, MA) and +2.5 kV compared to Earth ground. Peptide ions were surveyed ($MS^1$) in the orbitrap mass analyzer at 120,000 full width at half maximum (FWHM) resolution (AGC, $4 \times 10^5$ counts; maximum ion trap (IT), 50 ms) and fragmented by collision-induced dissociation (CID) using data-dependent acquisition as follows: high-pass threshold, $5 \times 10^3$ counts; dynamic exclusion, 30 s; mass tolerance, 10 ppm; monoisotopic peak determination, turned on; collision, in helium at 35% normalized collision energy (NCE). The generated fragment ions were detected in the ion trap analyzer using the following conditions: scan rate, turbo; AGC, $1 \times 10^4$ counts; maximum IT, 50 ms. The peptide ions were relatively quantified via multinotch $MS^3$ fragmentation of the top 10 $MS^2$ ions under higher-energy collisional dissociation (HCD) in nitrogen at 65% NCE. The $MS^3$ spectra were acquired at 50,000 FWMH in the orbitrap analyzer (AGC, $1 \times 10^5$ counts; maximum IT, 105 ms).

**Micro-LC MS Metabolomics.** A 2.5 µL of the metabolite extract was loaded onto a reversed-phase column featuring anion-exchange chemistry (Atlantis PREMIER BEH C18 AX Column, Waters, 1.7 µm, 2.1 × 100 mm) and separated at 500 μL/min in a micro-flow LC system (ACQUITY I-Class, Waters) using a gradient of buffer A (100% ACN) and buffer B (95% water, 5% ACN with 1 mM ammonium acetate; pH 7.8) as follows: 5% was held 0–0.5 min, then linearly increased to 90% over 9.5 min, and held at 90% for 3 min. To recover and recondition the analytical column before the next injection, solvent B was then decreased to 5% over 2 min and the column was rinsed for 7 min before the next injection. The column temperature was set to 30 ℃ during separation.

Metabolite ions were targeted in a trapped ion mobility time-of-flight mass spectrometer (timsTOF PRO, Bruker). The metabolites were ionized in a micro-flow electrospray source equipped with an ionBooster-ESI device with the following settings: end plate, 400 V; capillary voltage, 1,000 V; charge voltage, 300 V; vaporizer temperature, 240 °C; sheath gas, 1.5 L/min; nebulizer, 4.1 bar; dry gas, 2.6 L/min; dry temperature, 200 °C. Ions were detected in the negative ionization mode using the following settings: survey ($MS^1$) low- and middle-mass range, *m/z* 20–300 and *m/z* 50–1,300, respectively; spectral acquisition rate, 2 Hz. A list of metabolite ions was targeted using multiple reaction monitoring. Ions were isolated with 4-Da window (± 2 Da) for collision-induced dissociation (CID). The low-mass range (*m/z* 20–300) employed the following optimization for collision energy and *m/z* tuning: 12 eV at 115.0026; 12 eV at 145.0132; 12 eV at 168.9897; 12 eV at 184.9846; 10 eV at 191.0186. The middle-mass range (*m/z* 50–1,300) employed the following optimization for collision energy and *m/z* tuning: 15 eV at 338.9877; 18 eV at 346.0553; 25 eV at 426.0216; 20 eV at 505.9879.

**Adenylate Energy Charge Ratio.** Using external concentration calibration with authentic chemical standards, tissue-endogenous concentrations were determined for adenosine triphosphate (ATP), diphosphate (ADP), and monophosphate (AMP). Following the popular approach, we calculated the adenylate energy pool as the [ATP]/[ADP] ratio on the calculated concentration as well as the adenylate energy charge as follows: ([ATP]+0.5×[ADP]) / ([ATP]+[ADP]+[AMP]).

**GSH/GSSG Assay by Micro-LC MS.** A targeted assay was developed and optimized for GSH and GSSG as a quantitative measure of oxidative stress. A 2-µL volume of the metabolite extract was loaded onto an ACQUITY UPLC BEH amide column (130 A, 1.7 µm, 1 mm × 100 mm) and separated at 45 ℃ using a mobile phase delivered at 130 µL/min in a micro-LC system (Waters ACQUITY I-Class). The separation was performed in buffer A (water containing 0.1% v/v FA) with a gradient of buffer B (100% ACN with 0.1% v/v FA) as follows: 1% A for 0.05 min, then ramped to 80% A over 3.20 min, held at 80% A for 2 min, ramped to 1% A in 1.75 min. To recover and condition the analytical column for the next injection, buffer A was held at 1% for 7 min. GSH and GSSG were measured in the positive ion mode, and ionized in the ionBooster-ESI source, and detected on the timsTOF PRO (Bruker) with the following parameters: mass range, *m/z* 100–800; spectral acquisition rate, 4 Hz; scan mode, MRM (GSH: *m/z* 308.0911, charge state +1, retention time 2.5 min, width 4 Da (+/– 2 Da), and collision energy 26 eV); and GSSG (*m/z* 307.0833, charge state +2, retention time 2.9 min, width 4 Da (+/– 2 Da), collision energy 20 eV).

**GSH/GSSG Analysis by Glo Assay.** The SMO and RE tissues were dissected and snap-frozen on dry ice as described above. Each BR was prepared as a pool of 3 dissected tissues. The samples were extracted in 20% SS. The RE extract was diluted 20-times to ensure linearity for quantification based on prior validation using a concentration calibration curve. The GSH/GSSG-Glo Assay (Promega, Madison, WI) was used to assess GSH/GSSG ratios in the SMO and RE tissues. The assay was performed following vendor recommendations and tested for standards. The total GSH and GSSG levels were calculated based on a concentration calibration curve (0.5–50 μM of GSH). The GSH/GSSG ratios were compared between the SMO and the RE using a Student's t-test, after confirming data normality. A total of N = 5 BRs were analyzed for the SMO and the RE, each from the same clutch of embryos to reduce biological variability.

**$H_2O_2$ Imaging in Live Embryos**. The BioTracker Green $H_2O_2$ Live Cell Dye was reconstituted as per vendor recommendations and diluted to 10 μM in 20% Steinberg's solution. Embryos were obtained by natural mating and cultured to stage 9 after which they were incubated in the 10 μM BioTracker Green $H_2O_2$ Live Cell Dye at room temperature on a nutator until they developed to stage 10. The embryos were washed twice in 50% SS for 5 min each following staining and then imaged under brightfield and GFP channels using a stereomicroscope (SMZ18, Nikon Instruments Inc., Melville, NY). To assess differential enrichment of $H_2O_2$, the GFP fluorescence intensity was quantified in regions of the SMO as well as the dorsal and the vegetal portions of the stage 10 embryos that were contoured manually using the NIS-Elements software (Nikon). The measured intensity values were normalized to the background intensity, and differences in region-specific intensities were assessed using a paired Wilcoxon Signed Ranks Test (OriginPro version 2020b, Origin Labs, Northampton, MA).

**Injection of N-Acetyl Cysteine (NAC).** Using a microinjector (Warner Instruments), The SMO region of stage 10 embryos were microinjected with 4 nL of 0.5 M NAC (pH 7) as the experimental group or 20% Steinberg's solution (SS) as the control group. These experimental and control sibling embryos (wild-type, not experimented on) were cultured to stages 11 (mid-gastrulation) and 13 (neurulation) at room temperature for phenotype screening.

**Fibronectin Immunostaining**. NAC-injected and control siblings (injected with 20% SS), were fixed when controls reached stage 11 in 4% paraformaldehyde in MEM buffer (0.1 M MOPS, 0.5 M NaCl, 1 mM EGTA, 2 mM $MgSO_4$). The embryos were then washed and bisected along the midsagittal plane and processed for immunohistochemical detection of fibronectin as described elsewhere (10) using a mouse anti-fibronectin monoclonal antibody (1 μg/ml; Developmental Hybridoma Bank #4H2) and goat anti-mouse HRP-conjugated IgG (1:250, Cell Signaling #7076).

**Whole-Mount *In Situ* Hybridization**. The NAC-injected embryos were fixed in 4% paraformaldehyde in MEM (minimum essential buffer) at stages 11 and 13 and processed for whole-mount *in situ* hybridization as described elsewhere (11). Probes for *sox2*, *foxd4*, and *chd* were provided by Dr. Sally A. Moody (George Washington University, Washington, DC) and were prepared by *in vitro* transcription using Digoxigenin labeling mix (Roche).

**HRMS Data Analysis.** The proteomics and metabolomics data were processed following our established protocols. For <u>discovery quantitative proteomics</u>, the primary MS–$MS^n$ data were processed in Proteome Discoverer version 2.2 (Thermo) executing SEQUEST. The MS−MS/MS data were searched against the *Xenopus laevis* genome version *9.2* (downloaded from Xenbase on 12/06/2019) supplemented with the mRNA-derived PHROG version 1.0 database (12). The following search parameters were applied: enzyme, trypsin; number of missed cleavages, max 2; min number of unique peptides, 1; static modification, cysteine carbamidomethylation and lysine TMT6plex; variable modification, methionine oxidation, N-terminus acetylation; maximum mass deviation for main search of precursor masses, 10 ppm; *de novo* mass tolerance for tandem mass spectra, 0.6 Da; minimum score for modified peptides, 75. Peptide and protein identifications were filtered to <1% false discovery rate (FDR) against a reversed-sequence database. For targeted metabolomics, separate concentration calibration curves were experimentally determined to quantify selected endogenous metabolite concentrations in the SMO and RE. The metabolomics data were manually processed in DataAnalysis 4.0 (Bruker).

**Gene Naming and Canonical Annotation**. Gene Ontology (GO) annotation and enrichment analysis of gene classes and pathways were performed using the PANTHER database version 14.1 against *Xenopus tropicalis* as well as using Ingenuity Pathway Analysis (IPA, 2020 Qiagen) using human gene names for the identified proteins. Access to IPA was provided by Dr. Norman H. Lee (George Washington University, Washington DC). For convenience, both *Xenopus* and human gene names are provided in the **Electronic Supplementary Table 1** (**Table S1**). The Figures and text utilize human gene names, unless otherwise noted.

**Statistics.** Protein abundances were normalized based on total peptide signal in Proteome Discoverer 2.2. Protein identifications lacking reporter ion values were excluded from further analysis. Statistical data analysis was performed on log-transformed TMT abundance values. After validating the Null hypothesis, statistical significance was assessed in OriginPro 2020b using Student's t-test with a *p*-value < 0.05 marking significance. Data plots were created using OriginPro.

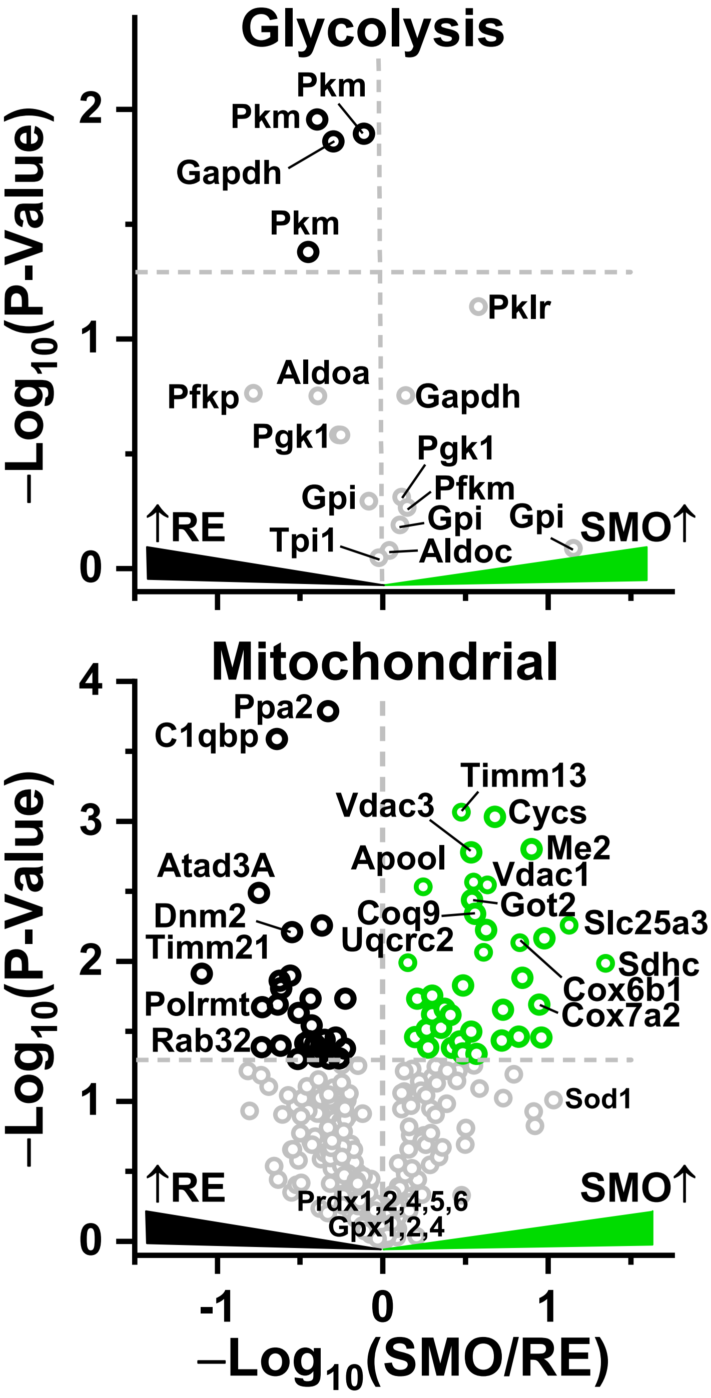

**Figure S1.** Quantitative comparison of glycolysis and mitochondrial proteins between the Spemann-Mangold Organizer (SMO) and the remainder of the embryo (RE). Key to statistics: Horizontal dashed line, statistical threshold at $p < 0.05$ (Student's t-test); Vertical dashed line, fold change = 1.

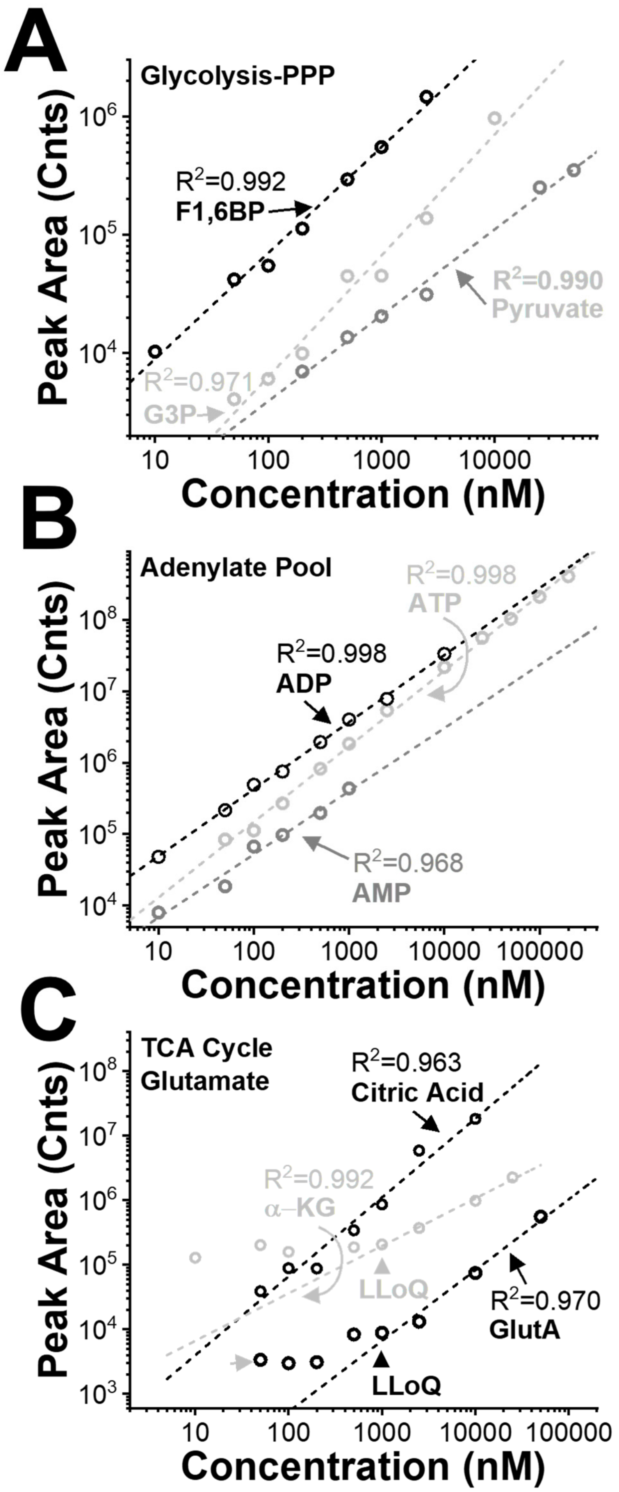

**Figure S2.** Metabolite concentration calibration curves for a panel of intermediates from (A) glycolysis, (B) phosphate energy pool (ATP, ADP, AMP), and (C) mitochondrial activity and including the TCA cycle. Key: R2, coefficient of linear regression.

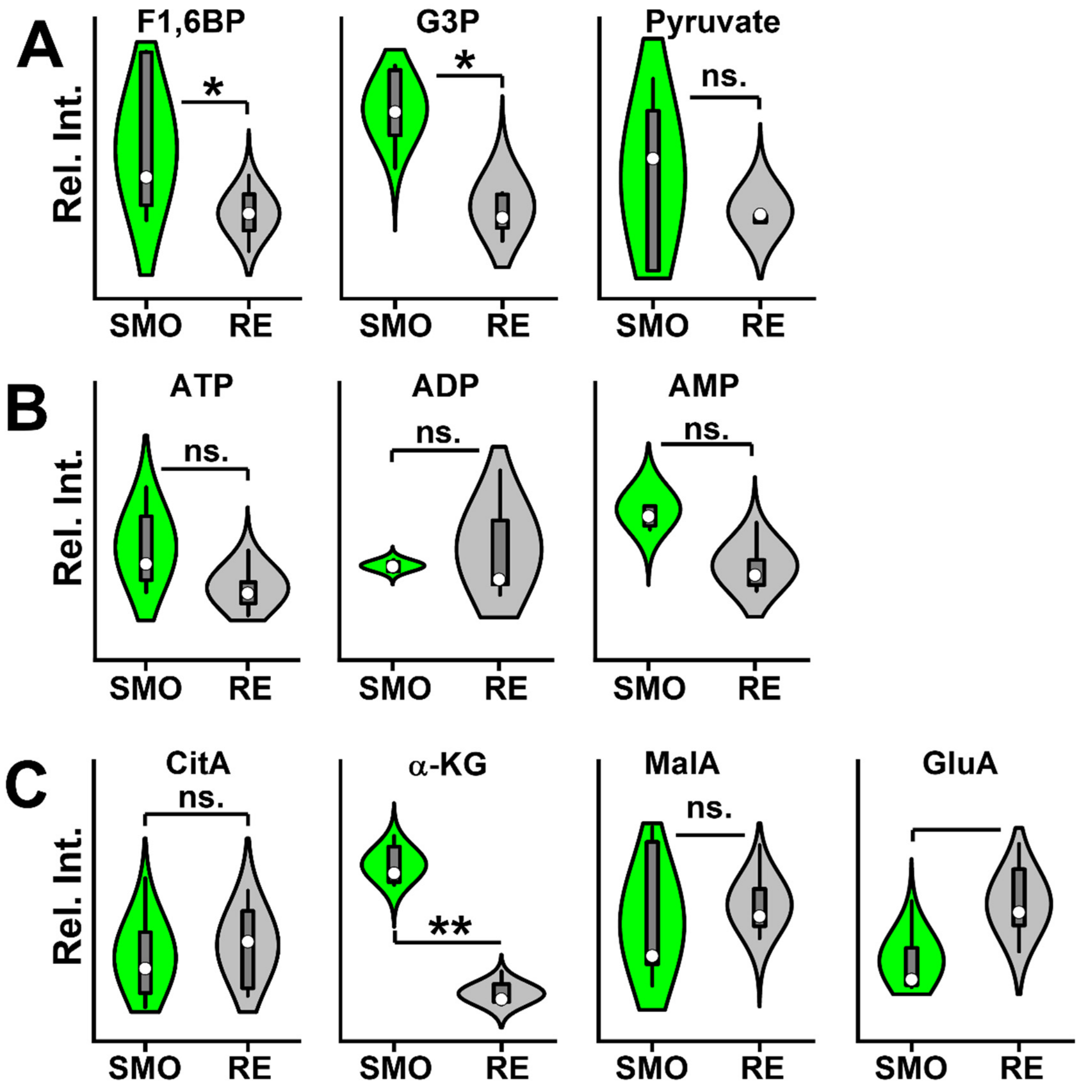

**Figure S3.** Metabolite profiling of the SMO and RE for targeted intermediates of (A) glycolysis, (B) the adenylate energy pool, and (C) mitochondrial activity including the TCA cycle. Key to metabolites: CitA, citric acid; F1,6BP, fructose-1,6-biphospate; G3P, glyceraldehyde-3-phospate; GluA, glutamic acid; MalA, malic acid; α-KG, α-ketoglutaric acid. Key to statistics: *$p < 0.05$; **$p < 0.005$ (Student's t-test).
